# Role of *Chymotrypsin-like Elastase 1* in Lung Physiology and in α1-Antitrypsin Deficiency

**DOI:** 10.1101/138776

**Authors:** Rashika Joshi, Andrea Heinz, Qiang Fan, Shuling Guo, Brett Monia, Christian E.H. Schmelzer, Anthony S. Weiss, Matthew Batie, Harikrishnan Parameswaran, Brian M. Varisco

## Abstract

α1-antitrypsin (AAT) deficiency-related emphysema is the fourth leading indication for lung transplantation. We previously demonstrated that AAT covalently neutralizes *chymotrypsin-like elastase 1* (*Cela1*) *in vitro*, that *Cela1* is expressed during the alveolar stage of lung development in association with regions of lung elastin remodeling, and that lung stretch increases *Cela1* expression and binding to lung elastin. Here we show that *Cela1* is exclusively responsible for stretch-inducible lung elastase activity, reduces postnatal lung elastance, and is required for emphysema in an antisense oligo model of AAT deficiency. *Cela1* mRNA is present in the human lung, and in the placental mammal lineage, *Cela1* is more conserved than *Cela2* or *Cela3* with unique promoter and protein elements indicating a unique role for *Cela1* in this lineage. These data demonstrate an adaptive role for *Cela1* in placental mammal lung biology with physiologic relevance to AAT-deficient lung disease in humans.

## Abstract

α1-antitrypsin related lung disease (AAT-RLD) is the fourth leading indication for lung transplantation and is characterized by protease-mediated progressive emphysema that manifests in the 4^th^ or 5^th^ decade of life. *Chymotrypsin-like elastase 1* (*Cela1*) is a digestive enzyme that binds to elastin in a stretch-dependent manner and is covalently neutralized by AAT. We hypothesized a role for *Cela1* in AAT-RLD. *Cela1*^-/-^ mice where phenotypically similar to wild type but had higher lung elastance and lacked stretch-inducible elastase activity. Wild-type mice administered anti-AAT oligo had reduced amounts of lung Cela1-AAT fusion protein in lung homogenate and spontaneously developed emphysema after 6 weeks. *Cela1*^-/-^ mice administered anti-AAT oligo were completely protected from these emphysematous changes. Cela1 recombinant protein did not require propeptide cleavage for elastolysis, and its elastolytic profile was similar to that of other pancreatic elastases. Phylogenetic analysis of vertebrate *Cela* promoter and protein sequences showed that placental mammal *Cela1* was distinct from other *Cela’s*, and that the placental mammal the *Cela1* gene was invariantly conserved despite variable loss of other *Cela* genes in non-carnivores. These data demonstrate that the pancreatic enzyme *Cela1* has been evolutionarily co-opted for a role in reducing lung elastance in the placental mammal lineage and that its stretch-regulated expression and elastolytic activity is responsible for emphysema in the absence of its anti-protease: AAT.

## BACKGROUND

Alpha-1-antitrypsin (AAT) neutralizes specific serine proteases in the lung and other tissues. Approximately 1:2000 individuals have AAT deficiency, and ten percent of these individuals will develop emphysema.^3^ AAT deficiency-related lung disease (AAT-RLD) is an underdiagnosed^4^ progressive emphysema that is the fourth leading indication for lung transplantation in adults.^5^ Unopposed neutrophil elastase, protease-3, cathepsin G, and matrix metalloproteinase activity have all been implicated in AAT-RLD pathogenesis;^6-8^ however, the standard of care, intravenous AAT-replacement therapy, only modestly slows emphysema progression in AAT-RLD.^9^ One major hurdle to understanding and treating AAT-RLD is the genetic heterogeneity and variable penetrance of the disease within different genotypes.^10^ Since the diagnosis of AAT deficiency is typically made only after the manifestation of disease symptoms,^11^ there exists an urgent need to understand the mechanisms underlying disease initiation and progression. The lack of a genuine animal model of AAT-RLD has hampered progress in this area.^12^

*Chymotrypsin-like elastase 1* (*Cela1*) is a digestive protease which is not expressed in the human pancreas,^13^ but which we and others have noted to be expressed in lung epithelial, intestinal, and immune cells.^1^,^14^,^15^ *Cela1* expression increases during post-pneumonectomy compensatory lung growth in a stretch-dependent manner,^2^ and Cela1-positive cells localize to regions of elastin remodeling during lung development.^1^ *Cela1* binds to lung elastin with stretch-dependent binding kinetics, and it is covalently bound by AAT *in vitro*^2^ as was previously reported for Cela2a.^16^Although these observations suggest that *Cela1* might play an important role in lung structure, function, and disease, this has yet to be tested in animal models.

To test whether Cela1 plays an important role in developmental and pathological lung remodeling processes, we created a *Cela1*^-/-^ mouse and expanded upon a previously published antisense oligo model of AAT deficiency.^17^ *Cela1*^-/-^ mice were viable and fertile with increased postnatal lung elastance and disordered lung elastin architecture. Anti-AAT oligo therapy induced emphysema from which *Cela1*^-/-^ mice were protected. *Cela1*^-/-^ mouse lung lacked the stretch-inducible lung elastase activity we previously described in wild type mouse lung.^18^ Although *Cela1* exhibited an elastolytic profile similar to that of other pancreatic elastases, it did not require propeptide cleavage for its elastolytic effect. Phylogenetic analysis of *Cela1* uncovered evidence that this gene has been uniquely conserved in the placental mammal lineage. Taken together, our data demonstrate that *Cela1* is a conserved gene which plays an important physiologic role during development and repair/regeneration but plays a maladaptive role in the absence of its cognate anti-protease.

## RESULTS

### Anti-AAT Oligo Model of AAT-RLD

To develop a model of AAT-RLD, we extended the duration of therapy of a previously published antisense oligo targeting all five murine AAT isoforms from three to six weeks.^17^ Once weekly administration of 100 mg/kg anti-AAT oligo resulted in a >99% decrease in liver AAT mRNA (the principal source of AAT, Figure 1A) and a 77.2% reduction in serum protein levels (p<0.001, n=8 per group, Mann-Whitney U-test) eight days after the 5^th^ dose of oligo. Since anti-sense oligo therapy can be associated with significant lung inflammation, we assessed inflammatory mRNAs in untreated, control oligo, and anti-AAT oligo treated lungs. There were no differences in the mRNAs of *macrophage chemoattractant protein-1* (*MCP1*, *CCL2*), *interleukin-6* (*IL6*), *interleukin-8* (*IL8*, *KC*, *CXCR1*), or *tumor necrosis factor alpha* (*TNFα*) in anti-AAT vs control oligo treated mice. Anti-AAT oligo treated mice had increased mRNA levels of the toll-like receptor 7 cytokines *interferon-alpha* and *beta* (*IFNα&β*); however, these levels were comparable to those of untreated 8-week mouse lung (Figure 1B). We also assessed the mRNA levels of proteases previously shown to be important in the pathogenesis of AAT-RLD. Anti-AAT oligo treatment did not increase the mRNA levels of *neutrophil elastase* (*ELANE*), *proteinase-3* (*Prot-3*), *cathepsin G* (*Cat G*), or *matrix metalloproteinases 12 & 14 (MMP12 & MMP14)*; however, *Cela1* mRNA was increased 8-fold (p<0.001, Figure 1C). There were no differences in weight gain between control and anti-AAT oligo treated mice (data not shown). Western blot of anti-AAT oligo treated lungs had a 77.3% reduction in the high molecular weight (∼70 kDa) AAT-Cela1 fusion protein that we previously reported (Figure 1D).^1^ These data validate our *in vitro* finding that Cela1 forms a covalent complex with AAT in the lung and support the hypothesis that *Cela1* is important in the pathogenesis of AAT-RLD.

**Figure 1.**
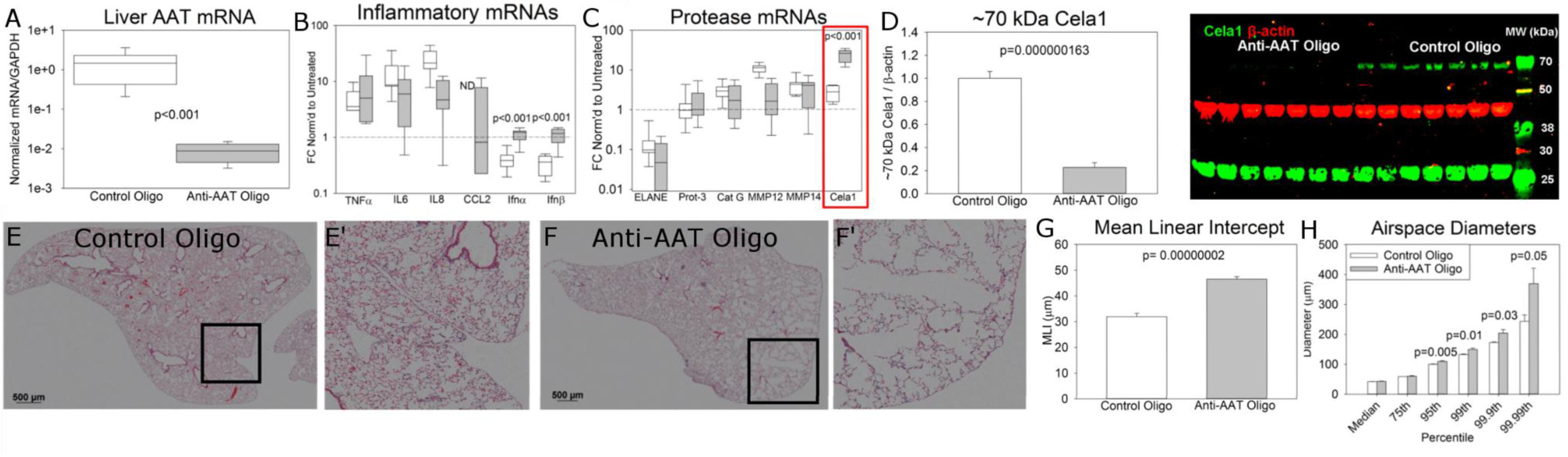
Mouse Model of AAT-RLD. **(A)** Anti-AAT oligo administration reduced liver AAT levels by >99% 8 days after administration of the final dose compared to control oligo. Note log scale and non-parametric data (n=8 per group). **(B)** Left lung inflammatory mRNAs were no different after treatment with anti-AAT or control oligo. Values were normalized to untreated adult control lung, and samples with undetectable mRNA were assigned a value of zero for statistical analysis. The *Interferon α&β* mRNA levels were elevated in anti-AAT treated lungs but comparable to untreated control lungs. (n=8 per group) **(C)** The mRNAs of many genes associated with AAT-RLD were no different in control and anti-AAT oligo treated lungs, but *Cela1* mRNA levels were increased 8-fold. **(D)** We previously reported a higher than expected molecular weight for Cela1 in mouse lung,^1^ and that Cela1 formed a covalent complex with AAT *in vitro*^2^ likely accounting for this observation. A 77% reduction in this high molecular weight form of Cela1 was observed in anti-AAT oligo treated lung compared to control-oligo treated lungs. Little difference was noted in the quantity of native molecular weight Cela1 however. n=7 per group **(E)** A representative middle lung lobe of wild type mouse treated with control oligo. Scale bar = 500 μm. **(F)** The emphysematous middle lobe of wild type mouse treated with anti-AAT oligo. **(G)** Mean linear intercept (MLI) values of WT mice treated with control or anti-AAT oligo (n=7 for control and 8 for anti-AAT). **(H)** While median and 75^th^ percentile alveolar diameters were no different in anti-AAT and control oligo treated lungs, diameters were significantly larger at percentiles at and above the 95th indicating that only the largest airspaces were contributing the increased MLI of anti-AAT oligo treated mice.

The lungs of anti-AAT treated mice demonstrated emphysematous changes reminiscent of the histologic changes of patients with AAT-RLD (Figure 1E-F). By the mean linear intercept method of airspace morphometry,^19^ the airspaces of anti-AAT oligo treated mice were 49% larger that control-oligo treated mice (Figure 1G). We further characterized lung morphometry my modifying a computerized morphometry program designed to quantify airspace heterogeneity in emphysema.^20^ This program uses local contrast to quantify airspace diameters of lung lobes from tile-scanned sections which are then used to create histograms (Supplemental Figure 1) from which descriptive statistics are derived. Using this technique, we determined that the difference in mean linear intercept was being driven by enlargement of airspaces in and above the 95^th^ percentile (i.e. airspace sizes were comparable below the 95th percentile, Figure 1H).

These data demonstrate that the murine anti-AAT oligo model of AAT-RLD faithfully phenocopies many elements of the human disease and suggest that AAT neutralization of Cela1 is important for preventing emphysema in the adult lung.

### *Cela1* Remodels Lung Elastin and Reduces Postnatal Lung Elastance

We created a global *Cela1*^-/-^ mouse on the C57 background using CRISPR-Cas9 gene editing and the guide RNAs indicated in Table S1.^21^ *Cela1* mRNA was absent from the pancreas of these mice, and there were no gross morphological differences or differences in weight gain between wild type and *Cela1*^-/-^ animals (Supplemental Figure 2). The lung architecture of *Cela1*^-/-^ animals at postnatal day (PND) 3, PND14, and PN 8 weeks was intact although airspaces were smaller in *Cela1*^-/-^ animals (Figure 2A, Supplemental Figure 2). Both the lung wet-to-dry weight ratio and percentages of cells from the distal lung were increased in *Cela1*^-/-^ mice (Supplemental Figure 2). Thus, *Cela1* did not appear to impact the alveolar septation program *per se*, but it did play a role in regulating distal airspace size.

**Figure 2.**
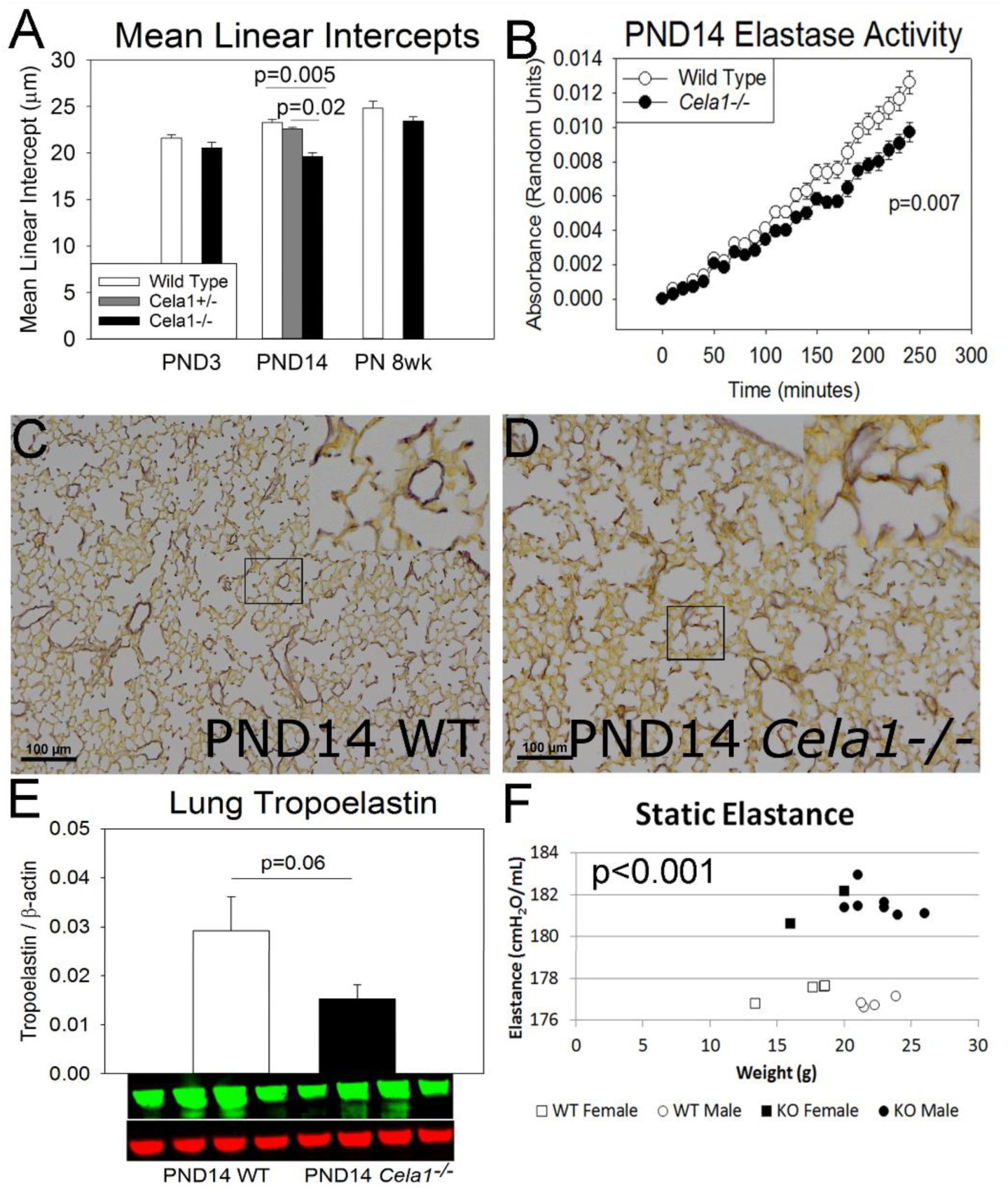
Impact of Cela1 on Distal Lung Structure and Function. **(A)** The mean linear intercepts of *Cela1*^-/-^ PND14 lungs (alveolar stage of lung development, n=9) were 18% smaller than wild type (n=6, *Cela1*^+/-^ n=5) with a trend towards smaller intercepts at PND3 (saccular stage of lung development, WT n=8, *Cela1*^-/-^ n=7) and PN 8 weeks (adult lung, WT & *Cela1*^-/-^ n=8). **(B)** The lung homogenates of *Cela1*^-/-^ PND14 mice had less elastase activity as compared to WT (n=6 per group). **(C)** PND14 WT lungs demonstrated the typical localization of dense elastin bands to septal tips. **(D)** *Cela1*^-/-^ lungs had less dense septal tip elastin bands and more diffuse elastin fibers distributed throughout distal airspace walls. **(E)** Western blot of PND14 lung homogenates indicated that *Cela1*^-/-^ mouse lungs had approximately half the soluble tropoelastin of WT lung. **(F)** As assessed by Flexivent, PN 8 week *Cela1*^-/-^ mouse lungs (n=9) were more elastic (i.e. more stiff) than wild-type (n=7) lungs. These differences were not age or sex dependent.

We next sought to assess the impact of *Cela1* on lung elastin and lung elastin remodeling. We previously reported that lung elastin remodeling is maximal during the alveolar stage of lung development.^18^ The elastase activity of *Cela1*^-/-^ lung homogenate was reduced by 25% compared to wild type at this stage (Figure 2B). Consistent with this finding, *Cela1*^-/-^ PND14 lungs had diffuse, thinner elastin fibers throughout the alveolar walls and lacked discrete elastin bands at the tips of secondary alveolar septae (Figure 2C-D). We therefore expected *Cela1*^-/-^ lungs to have more elastin than wild type; however, at PND14, *Cela1*^-/-^ lungs contained half as much tropoelastin as wild type by Western blot (Figure 2E). Morphometric analysis of elastin-stained PND3, PND14, and PN 8 week lungs confirmed this finding (Supplemental Figure 2). Furthermore, the mRNAs of all four elastin-associated genes assessed tended to be reduced in *Cela1*^-/-^ compared to wild type (Supplemental Figure 2). To assess the impact of these observed differences on lung dynamics, we measured lung elastance by flexivent in PN 8 week animals. *Cela1*^-/-^ mice had significantly higher elastance (lower compliance) than wild-type (Figure 2F).

The key findings from these experiments are that (1) *Cela1* contributes to lung elastin remodeling during development but this elastin remodeling does not impact overall lung elastin content, and (2) that *Cela1* reduces postnatal lung elastance which accounts for the reduced airspace size and expanded distal lung cell populations in *Cela1*^-/-^ mice. The finding that *Cela1*^-/-^ mice have less total lung elastin in the context of reduced lung elastase activity in combination with our previous report that Cela1 binds elastin with stretch-dependent kinetics suggest that Cela1 acts to remodel lung elastin in a localized, mechano-sensitive manner but that the principal determinant of postnatal lung elastin content is synthesis rather than remodeling with this synthesis being sensitive to local mechanical cues.^22,23^

### *Cela1* Is Required for Emphysema in the Anti-AAT Oligo Model of AAT-RLD

We previously demonstrated the covalent binding of bovine AAT (Serpina1) with murine Cela1 in the supernatant but not the cell lysate of *Cela1*-transfected cells.^2^ Since anti-AAT oligo treated mice had a reduction in the corresponding ∼70 kDa Cela1 band (Figure 1D), we treated *Cela1*^-/-^ mice with anti-AAT oligo to test whether *Cela1* was important in AAT-RLD pathogenesis. *Cela1*^-/-^ mice did not demonstrate any of the airspace destruction observed in wild type mice (Figure 3A compared to Figure 1G). The MLI of anti-AAT treated *Cela1*^-/-^ mice were significantly smaller than both anti-AAT and control oligo treated wild type mice (Figure 3B). Similarly, *Cela1*^-/-^ anti-AAT treated mice had smaller airspaces than control oligo treated wild-type mice at airspace diameters below the 95^th^ percentile. This difference is consistent with the previous observation of smaller distal airspaces in *Cela1*^-/-^ mice (Figure 2). More importantly however, *Cela1*^-/-^ mice demonstrated no increase in airspace size with anti-AAT oligo treatment (Figure 3C). The lack of emphysema in *Cela1*^-/-^ mice after administration of anti-AAT oligo supports a central role for *Cela1* in the emphysema of AAT-RLD.

**Figure 3.**
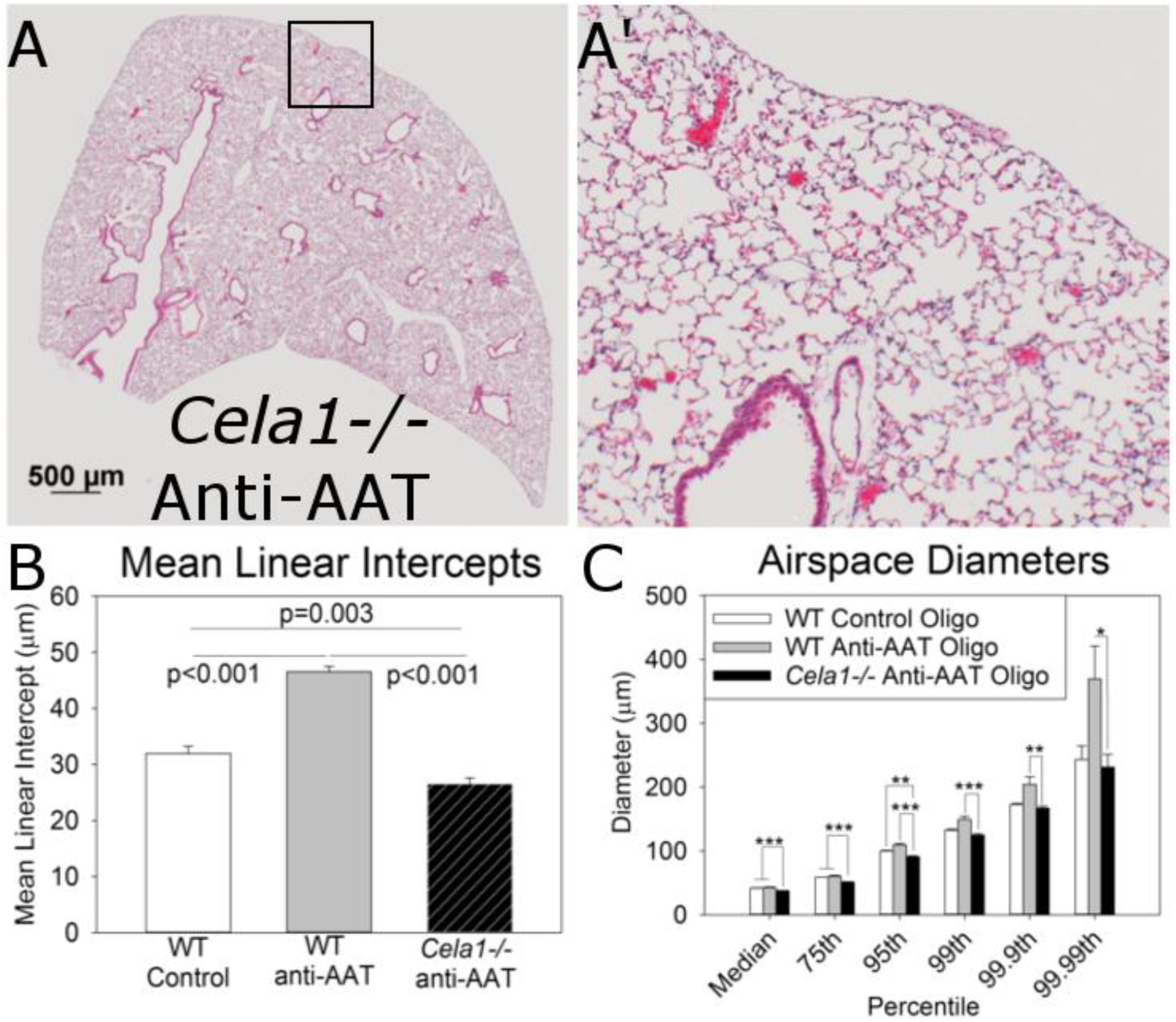
Requirement of Cela1 for Bullous Progression in AAT-RLD Model. **(A)** *Cela1-/-* mice treated with anti-AAT oligo for 6 weeks did not demonstrate any of the emphysematous changes seen in anti-AAT oligo treated wild type mice. Scale Bar = 500 μm. **(B)** As reported in Figure 1, wild type mice treated with anti-AAT oligo had increased MLI values, but the MLI values of *Cela1*^-/-^ mice (n=7) treated with anti-AAT oligo were smaller than WT. Consistent with data in Figure 2, the MLI of anti-AAT oligo treated *Cela1*^-/-^ mice were also smaller than control oligo treated wild type mice. **(C)** Comparing airspace diameter percentiles, the *Cela1*^-/-^ anti-AAT oligo treated mice had airspace diameter distribution comparable to wild type control oligo-treated mice. *p<0.05, **p<0.01, ***p<0.001.

### *Cela1* in Human Lung

To demonstrate the potential human health significance of Cela1-AAT interaction, human lung tissue was obtained from an organ donor to assess for *Cela1* expression. The presence of *Cela1* mRNA was confirmed by sequencing of PCR products (Supplemental Figure 3); however, neither our guinea pig or commercially available rabbit Cela1 antibodies were unable to detect the protein by Western blot. While we could not prove the protein-level expression of Cela1 in human lung, Cela1 mRNA is present in the human lung.

### Characterization of the Cela1 Elastolytic Profile

Since we previously demonstrated that Cela1 binds to lung elastin with stretch-dependent binding kinetics,^2^ we wished to more fully characterize the elastolytic profile of *Cela1*. To determine the importance of Cela1 in the previously described stretch-dependent lung elastase activity,^18^ we utilized a biaxial lung stretching elastin device (Supplemental Figure 4) with fluorescent elastin *in situ* zymography using the lungs of *Cela1*^-/-^ and wild type mice with and without stretch. As previously demonstrated, the elastase signal of wild type lung did not increase without stretch, but there was increased elastase activity with sequential increases in lung stretch. *Cela1*^-/-^ mice lacked this stretch-inducible elastase activity (Figure 4A-D). These data indicate that Cela1 is entirely responsible for stretch-inducible elastase activity in this *ex vivo* murine lung model.

**Figure 4.**
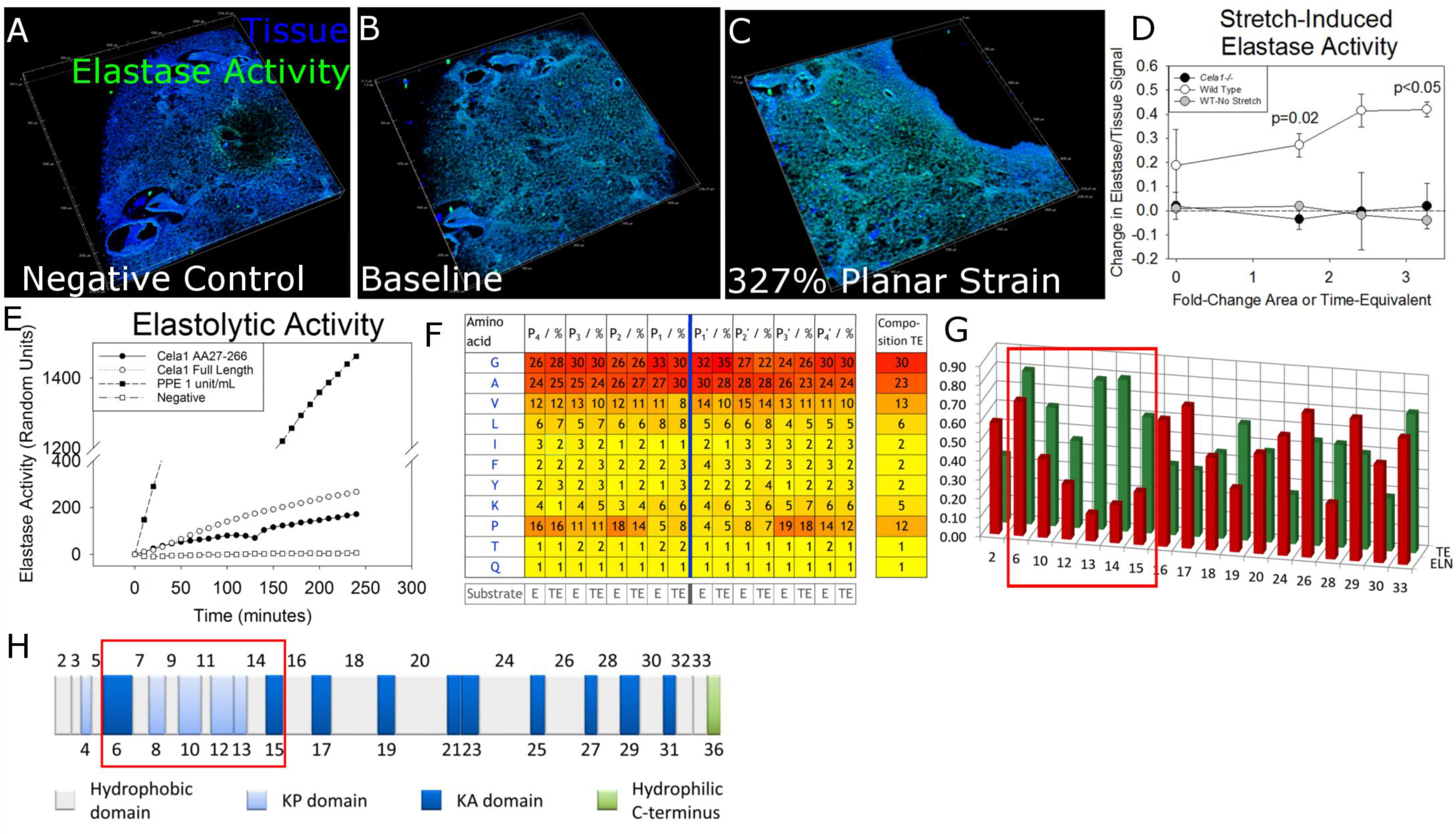
Elastolytic profile of Cela1. **(A)**The lungs of 12 week-old mice were inflated with gelatin, 200 μm sections were adhered to the silicone insert of a 3D-printed confocal lung stretching device, and the gelatin was removed. Lung tissue was defined by autofluorescence, and elastase activity determined using a soluble elastin substrate conjugated to a quenched fluorophore. The 3D reconstruction of a lung section is shown with each major tick mark representing 500 μm. **(B)** After application of the zymography substrate, there was an appreciable increase in the elastase signal. A representative wild type lung section is shown. **(C)** The same lung section is shown at the maximal strain achievable with this device. An increase in the elastase signal is appreciable. (D) The stretch-induced lung elastase activity of wild type (n=3) and *Cela1*^-/-^ (n=2) lung sections was compared. Wild type lung not exposed to stretch but incubated with substrate for an equivalent period of time did not demonstrate an increase in elastase signal. *Cela1*^-/-^ lung which was stretched lacked the inducible elastase activity that was observed in wild type lung. Comparisons are by one way ANOVA. **(E)** As assessed using a plate-based fluorometric elastase assay, full-length Cela1 demonstrated elastolytic activity comparable to that of Cela1 without signaling and propeptide. **(F)** Full length, recombinant Cela1 was incubated with soluble human tropoelastin isolated human skin elastin and degradation products were analyzed. While Cela1 had a propensity for hydrophobic residues in its binding pocket, the only amino acid that was non-preferred in regions adjacent to cleavage sites was proline. However, proline was preferred at P_4_-P_2_ and P_3_’-P_4_’ sites suggesting that the conformational turn induced by proline residues at these locations enhanced substrate binding. **(G)** While there was no major difference in the amino acid residues cleaved by Cela1, mature elastin had substantially fewer cleavage sites in domains 6-15 (red box). **(H)** While there was no major difference in the hydrophobicity of these domains (ivory color, domains numbered above simplified tropoelastin protein sequence) compared to others, these domains contain crosslinking sites (blue color). Since soluble tropoelastin lacks these crosslinks, they likely account for the difference in elastin degradation between mature elastin and soluble tropoelastin.

*Cela* family members and related *Chymotrypsin* proteases are synthesized as zymogens and activated in the intestinal lumen after interaction with pepsin. However, the protease that would accomplish this task in the lung was unclear. We therefore synthesized recombinant full-length Cela1 in a eukaryotic system and compared its elastolytic activity with previously published truncated Cela1 (without propeptide) synthesized in a prokaryotic system.^2^ Full-length Cela1 had a level of elastolytic activity comparable to the truncated Cela1 (Figure 4E). Thus, Cela1 does not require propeptide cleave for its elastolytic activity.

Lastly, we wished to understand the specificity with which Cela1 cleaved tropoelastin and elastin. In the lung and other elastin-containing tissues, translated tropoelastin forms coacervate droplets due to its high number of hydrophobic domains. These coacervates are directed to microfibrils and formed into elastin fibers coordinately with a range of microfibril-associated proteins.^24^Tropoelastin monomers on adjacent microfibrils are crosslinked at lysine residues after oxidative deamination by lysyl oxidase and related proteins increasing fiber strength.^25^ Porcine pancreatic elastase, which is largely composed of *Cela*-family members, cleaves elastin and tropoelastin in a relatively aggressive and nondiscriminatory manner.^26^ To determine whether full-length Cela1 possesses a similar elastolytic pattern, we performed a degradation analysis of recombinant human tropoelastin and mature human elastin. Full-length Cela1 degraded both elastin and tropoelastin aggressively and without substantial site specificity-the only specificity being a preference for prolines outside of the P_1_-P_2_’ positions (Figure 4F & Supplemental Figure 5). However, there was a substantial difference in tropoelastin and elastin degradation with regards to the domains cleaved. While the fraction of amino acids with documented cleavage sites was relatively consistent across tropoelastin domains, cleavage of domains 6-15 was reduced in elastin compared to tropoelastin (Figure 4G). These domains involve or are adjacent to lysine- and proline-containing (KP) crosslinking domains 4, 8, 10, 12, and 13 (Figure 4H) which most likely deny Cela1 substrate access accounting for the differentially degradation of mature elastin and recombinant tropoelastin. The elastolytic specificity of *Cela1* is similar to that of other pancreatic elastases,^26^ and the elastolytic activity of full-length Cela1 is regulated by steric hindrance of crosslinking residues.

Taken together, these findings indicate that Cela1 is not a zymogen and that its elastolytic profile is similar to that of other pancreatic elastase being constrained only by the conformational changes of proline residues and by steric hindrance of crosslinking domains. Cela1 is responsible for stretch-induced elastase activity in lung, although the exact mechanisms are unclear.

### Evolutionarily Conserved Role for Cela1 in Reducing Distal Lung Elastance in Placental Mammals

Given our findings that *Cela1* was expressed in the lung and played a role in reducing postnatal lung compliance, we wondered whether the lung expression of *Cela1*, which here-to-for has been assumed to only be involved in digestive processes, could have been conferred an evolutionary advantage to groups of animals which require expansion of air-exchanging surfaces for respiration. Mammals respire differently than other vertebrate clades. Reptiles and birds respire by using abdominal and thoracic muscles into air sacs with subsequent passage through gas-exchanging respiratory canals.^27^ Mammals, in contrast, have a muscularized diaphragm and respire by diaphragmatic contraction which increases the volume of the gas-exchanging units reducing the transpulmonary pressure gradient with subsequent ingress of air. High elastance lungs would impede this process increasing the work of breathing, and processes to reduce lung elastance might provide an evolutionary advantage.

To define the evolutionary conservation of *Cela1*, we used all available Ensembl sequences to construct phylogenetic trees of the *Cela* promoter and protein sequences. While mammalian *Cela2* and *Cela3* promoter sequences were interspersed within the different non-mammalian *Cela* promoter sequences, mammalian *Cela1* promoter sequences were phylogenetically distinct (Figure 5A). Similarly, mammalian Cela1 protein sequences were phylogenetically unique from mammalian Cela2, Cela3, and non-mammalian Cela sequences (Figure 5B). In comparing the two phylogenetic trees, the only mammals with *Cela1* protein and promoter sequences distant from the other mammalian *Cela1* sequences were the platypus (*Ornithorhynchus anatinus*) and Tasmanian devil (*Sarcophilus harrisii*). Thus, it seems that *Cela1* promoter and protein sequences have been uniquely conserved compared to the other *Cela* isoforms over the course of placental mammal evolution.

**Figure 5.**
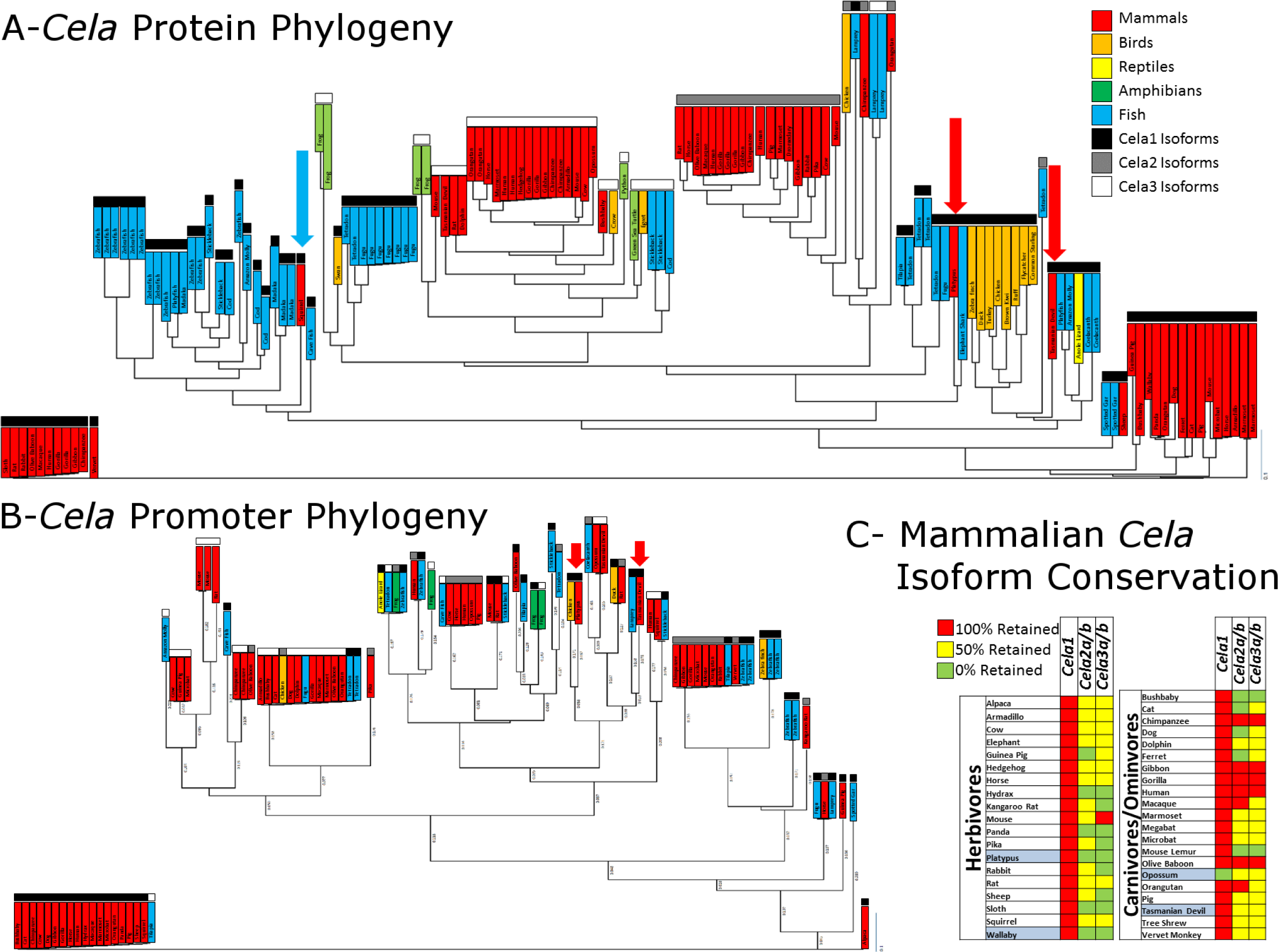
Unique Role for *Cela1* in the Placental Mammal Lineage. **(A)** Cela1 protein sequences were analyzed with clustering of mammalian Cela1 protein sequences away from mammalian Cela2&3 and non-mammalian Cela1, 2, and 3. Common names are listed. Different vertebrate clades are color coded, and sequences for *Cela1, 2, & 3* isoforms are coded in black, grey, and white respectively. Only two mammalian Cela1 protein and promoter sequences failed to cluster with the other mammalian Cela1 sequences: the platypus and the Tasmanian devil (red arrows). The rabbit Cela1 protein sequence included introns, but is reported for completeness (blue arrow). **(B)** Phylogenetic tree of Ensembl 200 base pair promoter sequences for all *Cela* family members. Like the pattern observed with protein sequences mammalian *Cela2&3* promoter sequences were similar to non-mammalian *Cela1, 2, & 3* sequences, mammalian *Cela1* sequences were less similar. Again, the Tasmanian devil and platypus *Cela1* sequences failed to cluster with other mammalian *Cela1* sequences. **(C)** Using Ensembl and Entrez Gene, the absence or presence of the *Cela1* gene (green or red respectively), or presence of zero, one, or two *Cela2* and *Cela3* genes (red, yellow, and green respectively) were plotted in a simplified heat map. Annotated mammals were segregated by diet, and non-placental mammals were highlighted in blue. Except for the opossum, all annotated mammals had retention the *Cela1* gene while herbivores had more frequent loss of *Cela2* and *Cela3* isoforms. Primates had higher retention of both *Cela2* and *Cela3* isoforms.

In evolutionary biology, duplicated genes that are not adaptive are lost over time.^28^ To determine whether there were any differences in the retention of *Cela1*, *Cela2a&2b*, and *Cela3a&3b* isoforms over the course of mammalian evolution, we used Ensembl and Entrez Gene to identify which genes were retained in the genomes of all annotated mammalian species. *Cela1* was retained in 39 of 40 mammals (being absent only in opossum, *Monodelphis domestica*, a marsupial). Ten of 19 herbivores had loss of all *Cela2&3* isoforms whereas only 2 of 21 carnivores/omnivores demonstrated such loss (Figure 5C). These two omnivores were the bushbaby (*Otoemur garnettii*) and mouse lemur (*Microcebus murinus*), both of whose diet consists primarily of fruits and insects suggesting loss of an unused gene. Most primates had retention of all four *Cela2&3* isoforms. These data suggest that while *Cela2&3* redundantly fulfill a digestive role and have been variably lost in mammals that do not consume elastin-containing foods, *Cela1* has been retained supporting the assertion that *Cela1* plays a unique biological role in placental mammal biology.

## DISCUSSION

We have shown that *Cela1* is a uniquely conserved lung protease that reduces postnatal lung elastance, and that AAT neutralization of Cela1 is required to preserve airspace architecture in the adult lung. While the expression of *Cela1* during lung development,^1,14^ and post-pneumonectomy compensatory lung growth^2^ has been reported, this is the first report defining its role in postnatal lung morphogenesis, physiology and pathophysiology. Although other proteases such as *neutrophil elastase*, *protease-3*, *cathepsin G*, and *matrix metalloproteinases 12* and *14* ^7,29-31^ have been implicated in the pathogenesis of AAT-deficient emphysema, this is the first report of a role for *Cela1* in this disease and the first report implicating one of these proteases in a model that specifically reduces serum levels of AAT. Thus, we have identified a targetable protease that is critical for postnatal lung physiology and is critical in the pathobiology of AAT-RLD.

Our work highlights how evolutionary concepts can lead to a greater understand developmental and disease processes. Following gene duplication events, daughter genes undergo synonymous and nonsynonymous substitutions at a five-fold greater rate than parent genes.^28^ If daughter genes fulfill an adaptive role, then they are retained, and if not, they are lost. In the case of *Cela1*, this gene was retained in placental mammals while other *Cela* genes were lost in species that do not ingest elastin. By reducing the work of breathing, the adaptation of *Cela1* for a role in respiratory physiology likely conferred a selective advantage to the placental mammal lineage. Since *AAT* is more ancient than *Cela1*,^32^ and other Cela isoforms are not neutralized by AAT,^16^ it is likely that the amino acid substitutions that differentiate placental mammal *Cela1* from other placental mammal *Cela* isoforms and from all other vertebrate *Cela’s* are responsible for this adaptive and functional difference. However, this evolutionary, physiologic adaptation becomes a driver of disease pathology in the absence of *Cela1’s* anti-protease: AAT.

In summary, we have shown that *Cela1*, a digestive protease, has been adapted to fulfill important role in mammalian lung matrix remodeling, but that in the absence of its cognate anti-protease, AAT, its remodeling activity is unchecked leading to emphysema.

## ACKNOWLEDGEMENTS

We would like to acknowledge the grant funding which made this project possible

- Cincinnati Children’s Research Foundation (CCRF) Procter Fellowship Award (BMV)
- Parker B. Francis Fellowship Award (BMV)
- NHLBI K08HL131261 (BMV)
- German Research Foundation (DFG) Grant HE 6190/1-2 (AH)
- Fraunhofer Internal Program Grant Number Attract 069-608203 (CEHS)

We also acknowledge the following individuals for their assistance with this project

- John Matthew Kofron and the CCRF Confocal Imaging Core for assistance with image acquisition and analysis
- Monica Delay and the CCRF Flow Cytometry Core for assistance with flow cytometry experiment design and analysis.
- Xie Hurang, formerly of the CCRF Mouse Transgenic Core and not at the Michigan State Mouse Transgenic Core for assistance with CRISPR guide RNA design, validation, and *Cela1*^-/-^ mouse creation.
- Satish Madala of the Cincinnati Children’s Hospital Medical Center Division of Pulmonary Medicine provided assistance with Flexivent experiments.
- Patrick Lahni of Cincinnati Children’s Hospital Division of Critical Care Medicine assisted with running image analysis programs on an institutional computing cluster.

## AUTHOR CONTRIBUTIONS

- RJ created *Cela1*^-/-^ mice, designed and performed most animal and lung histology experiments
- AH performed elastin degradation analysis and provided insight into the nature of Cela-mediated proteolysis
- QF identified gain of function in full-length Cela1 and improved upon live lung sectioning technique
- SG performed validation and preliminary experiments of anti-AAT and control oligos
- BM developed the anti-sense oligo therapy used in AAT silencing
- CEHS designed mass spectrometric experiments and analyzed elastin degradation data
- MB designed and constructed the 3D-printed confocal lung stretching device
- ASW identified connection between crosslinking domains and differential degradation of soluble and mature elastin
- HP wrote the MatLab airspace morphometry program
- BMV-Identified the evolutionary relationship between Cela isoforms and conceptualized, coordinated, and executed the project.

## Competing intests

BMV and CCHMC hold a provisional patent on the use of anti-Cela1 silencing strategies for the treatment of AAT-RLD.

SG and BM are employees and stock holders of Ionis Pharmaceuticals.

## Supplementary Data

**Table S1.**
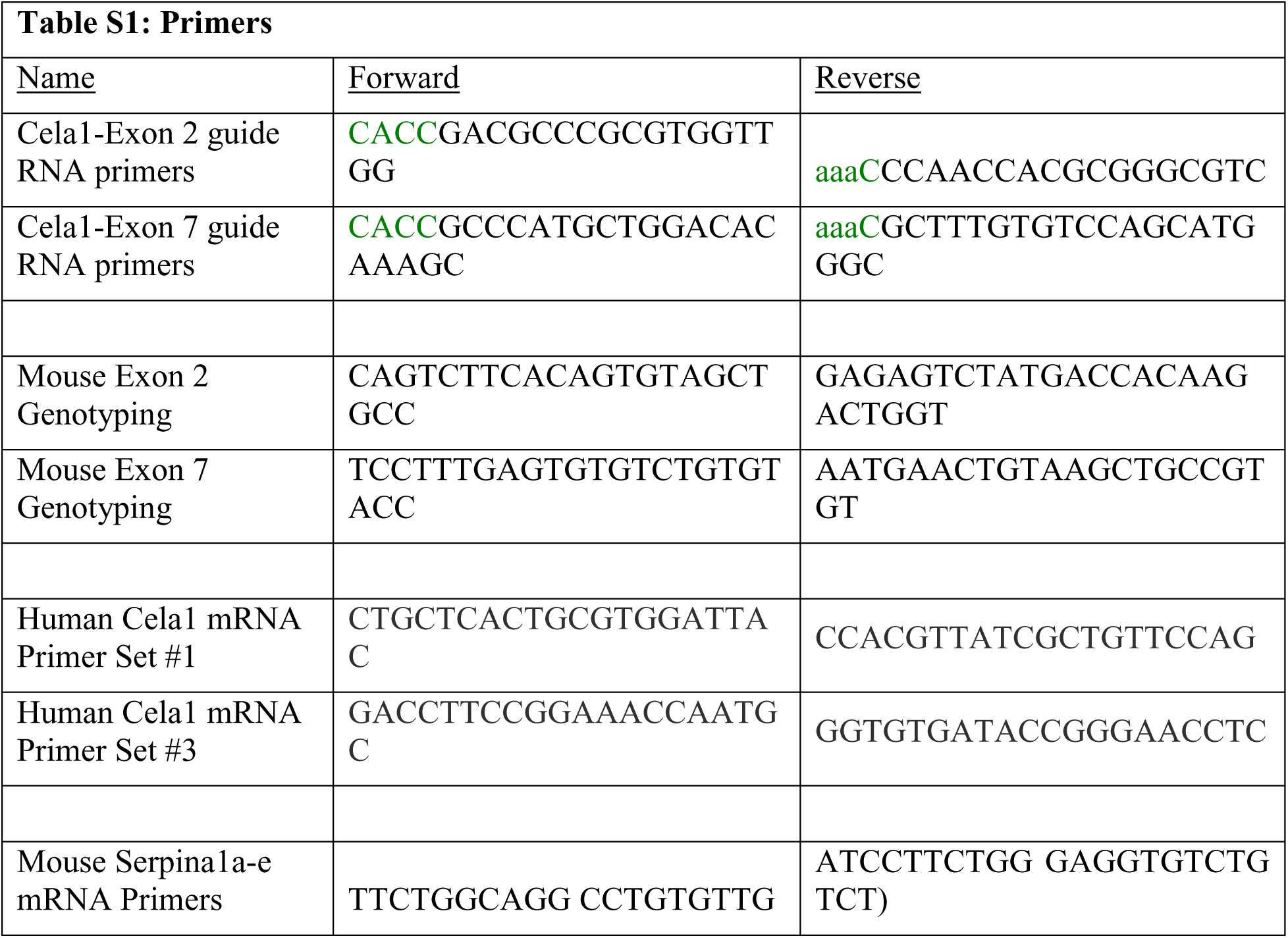
Primers Used.

**Supplemental Figure 1.**
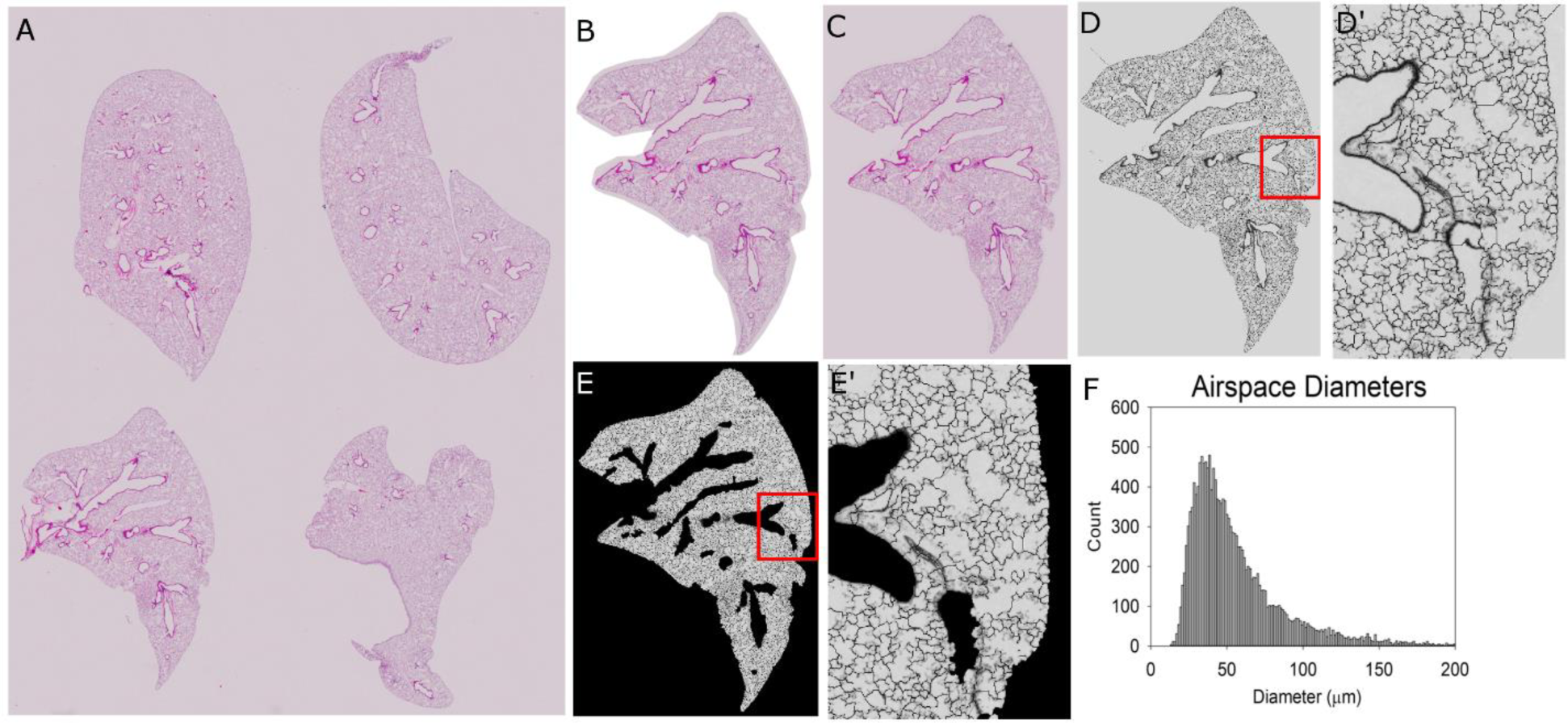
Method for Automated Airspace Diameter Measurements. **(A)** After fixation, lung lobes were disarticulated, paraffinized and arranged in a consistent pattern during embedding. Five μm sections were tile scanned at 4X magnification. **(B)** Each lobe was cropped using GIMP, and (C) pasted onto a new canvas matching the background color. **(D)** Airspace segmentation was achieved using the noted MatLab program. **(E)** Background, major vessels, and major airways were excluded from analysis using the “blackout” MatLab Program which also outputs all diameters. **(F)** Histograms were created from these diameters and descriptive statistics used for comparative analysis.

**Supplemental Figure 2.**
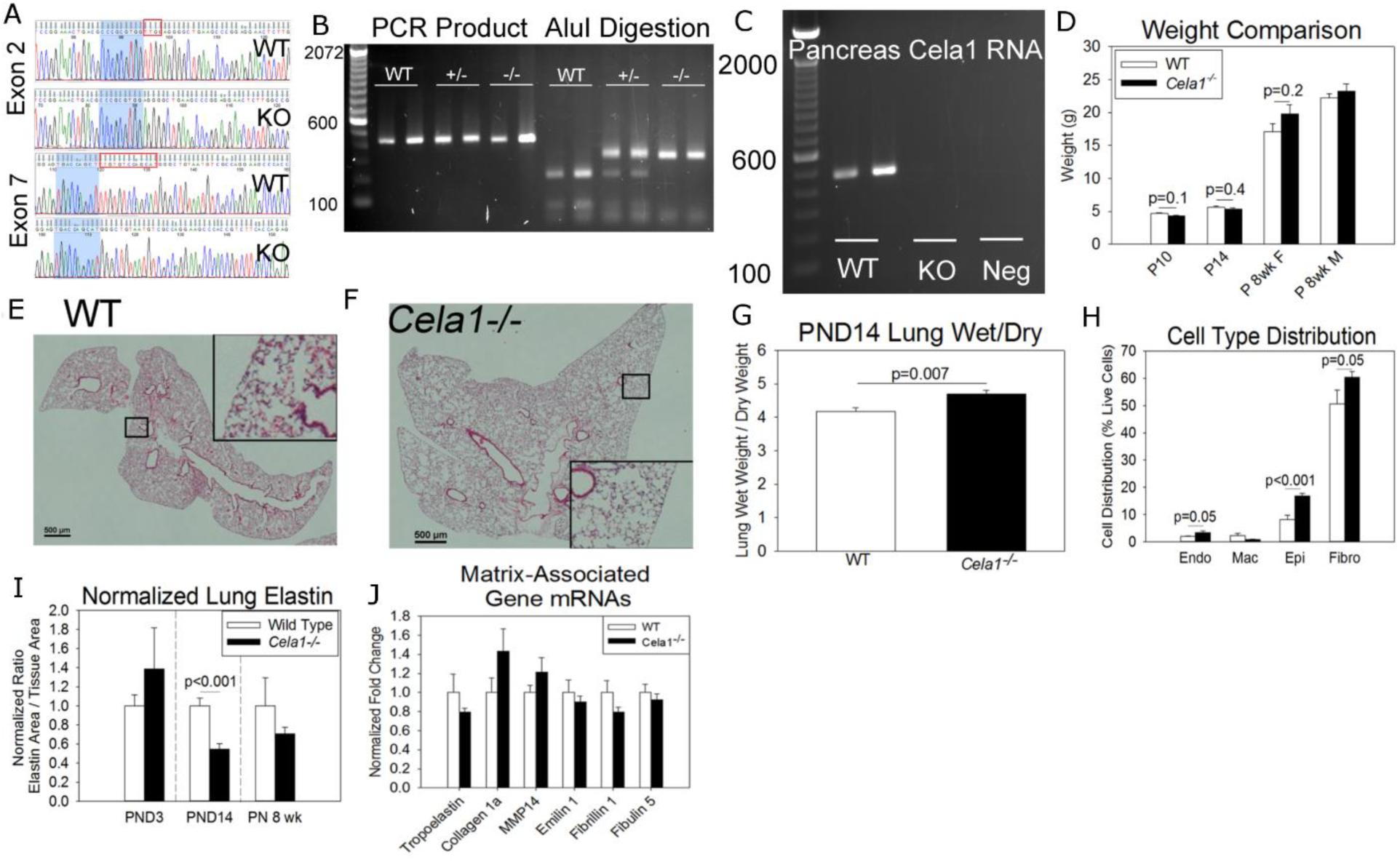
The *Cela1-/-* Mouse. **(A)** Cela1 Exon-2 and Exon-7 guide RNAs (Table S1) were injected into fifty single cell C57BL/6 mouse embryos which yielded a single mouse with a four base pair deletion in exon-2 and a twelve base pair deletion in exon-7 (indicated by red boxes). These deletions were determined to be on the same allele being co-segregated in three generations of heterozygous and homozygous mice. **(B)** Genotyping PCR of Cela1 exon 7 demonstrated the near-equivalence of PCR product weights between wild type, *Cela1*^+/-^, and *Cela1*^-/-^ mice. To overcome this limitation, the removal of an AluI endonuclease site was used to differentiate between genotypes. **(C)** PCR of adult mouse pancreas demonstrating the absence of Cela1 mRNA in *Cela1*^-/-^ mice. **(D)** The body weight of wild type and *Cela1*^-/-^ mice at different ages was comparable. **(E)** At postnatal week 8 (PN 8 wks), there was no appreciable difference in distal lung morphology between WT and **(F)** *Cela1*^-/-^ lungs. Accessory lobes are compared. Scale bar = 500 μm. **(G)** PND14 *Cela1*^-/-^ lungs had a higher wet to dry lung weight ratio indicating greater tissue content (n=6 per group). **(H)** Flow cytometry of PND 14 lungs demonstrated increased relative abundances of endothelial cells, epithelial cells, and fibroblasts, n=6 per group. **(I)** Morphometric quantification of total lung elastin demonstrated a 45% reduction in lung elastin at PND14 and a 29% reduction at PN 8 weeks. To account for differences in lung density, values were normalized to tissue area, but similar findings were present when normalized to lung area. **(J)** In PND14 lungs, differences in lung matrix-associated mRNAs were modest, but all assayed elastin-associated genes were reduced in *Cela1*^-/-^ compared to WT (n=6 per group).

**Supplemental Figure 3.**
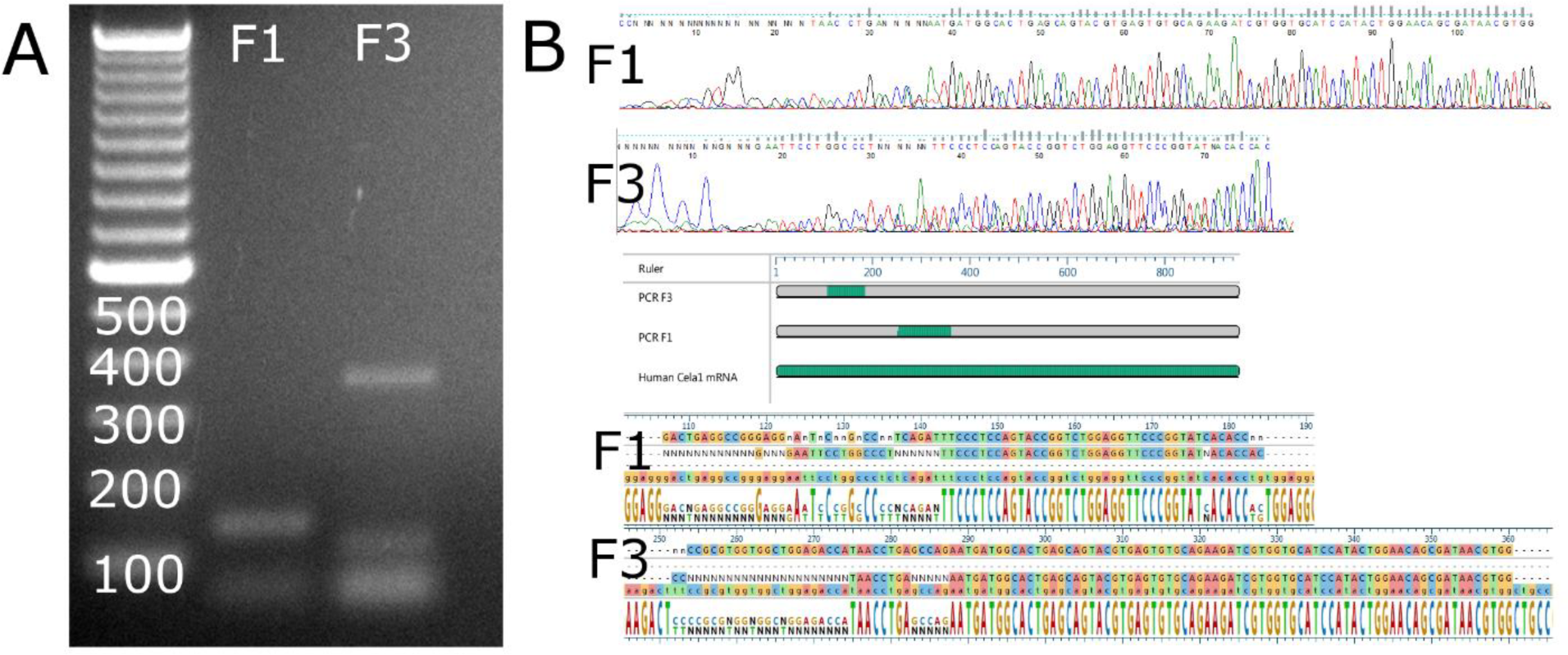
Presence of Cela1 mRNA in Human Lung. (A) Two different primer sets for *Cela1* mRNA displayed bands at the predicted weights (ladder is in base pairs). (B) Sequencing the F1 and F3 PCR products yielded satisfactory sequences as evidenced by the chromogram images. Alignment of these sequences with human Cela1 mRNA mapped to the predicted locations. Sequence alignment showed good agreement.

**Supplemental Figure 4.**
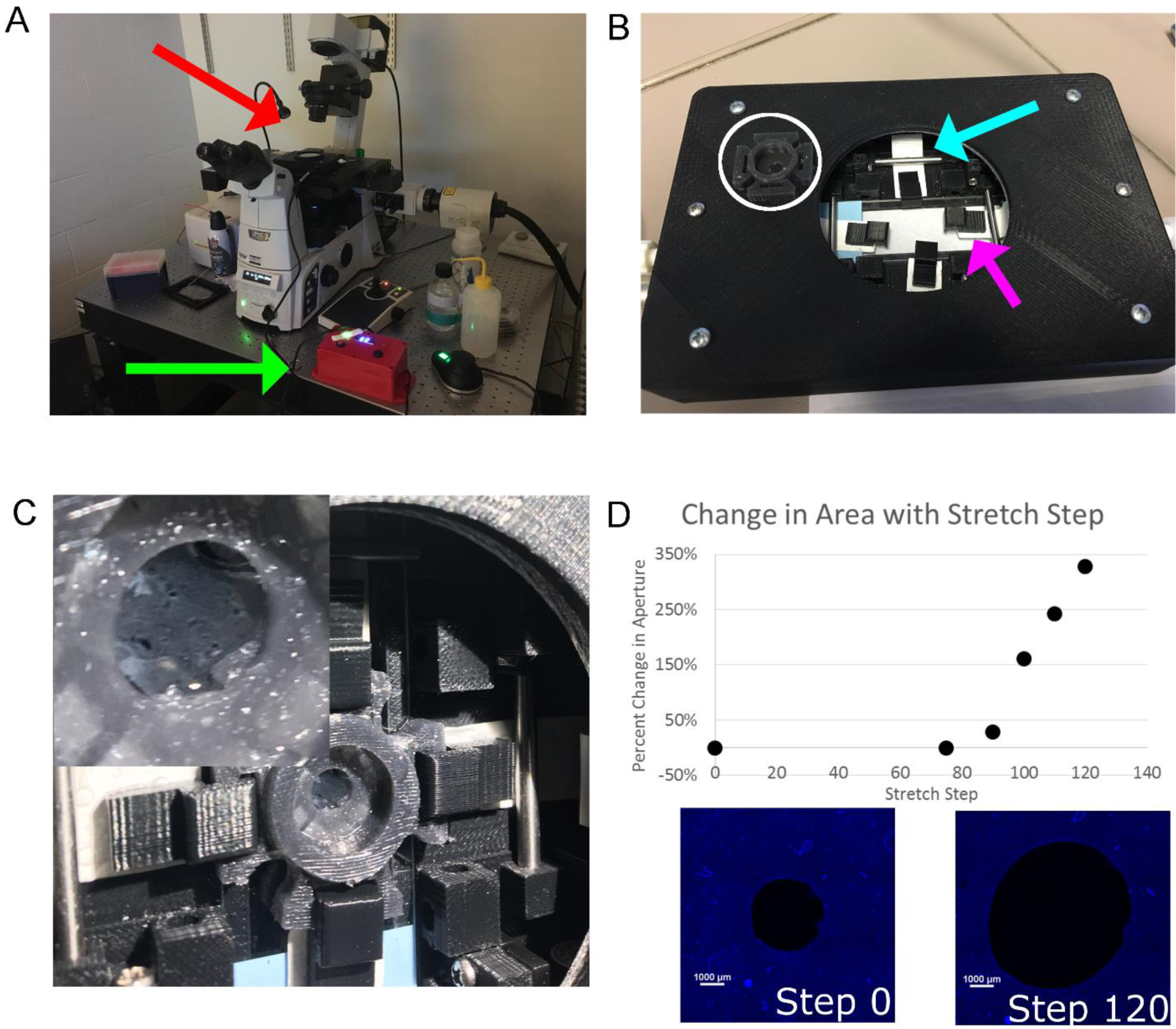
3D-printed Confocal Stretching Device. (A) The lung stretching device (red arrow) sat within a K-mount of an inverted Nikon A1 LUN-V microscope. The stretching device was connected to a control box (green arrow) which advanced the servo motors within the lung stretching device. (B) Four servo motors were housed within a custom 3D-printed case. Each motor turned an axle which is attached to Tyvek strips (aqua arrow) with 3D-printed clips (magenta arrow) at the end. A polysine slide sat below the strips and rolling bars keep the Tyvek strips and plastic clips in a stable Z-plane. Lung sections were adhered to a silicone mold (white circle) at four points using acrylic glue and the inflating gelatin media was removed as described in methods. (C) The silicone well was secured to the stretching device with clips. A stretched lung section is shown inset. (D) Chart of how different steps of stretch increased the mold aperture. Note that 75 steps were required to relieve the slack in the system. The apertures at step zero and step 120 are shown with a 1000 μm scale bar.

**Supplemental Figure 5.**
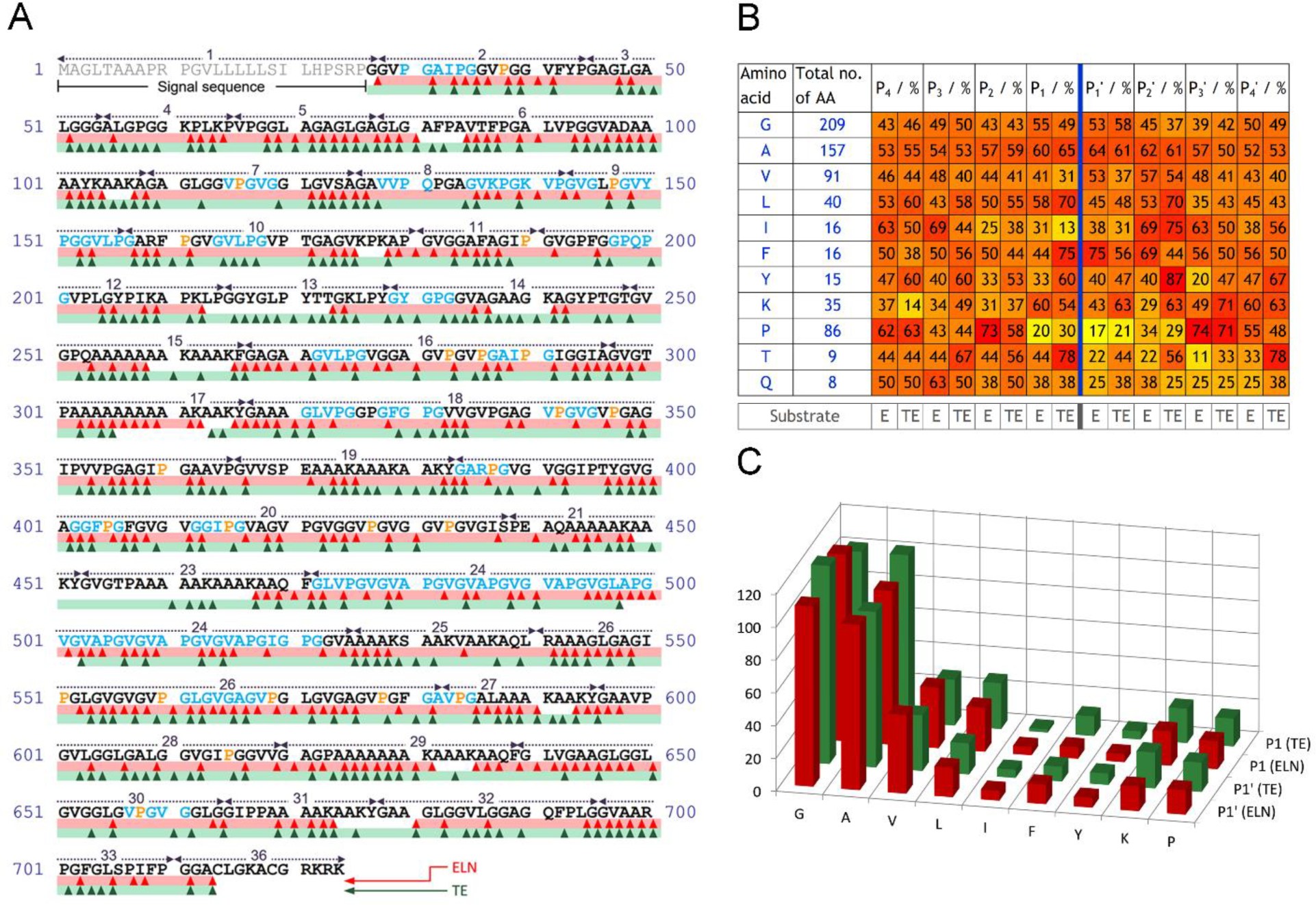
Cela1 Degradation of Elastin. (A) An elastolysis map of soluble tropoelastin (TE, red) and mature elastin (ELN, green) incubated with Cela1 demonstrating the amino acid residues at which each molecule was cleaved. Domain numbers are indicated above the protein sequence of human tropoelastin isoform 2. (B) Table of the frequency with which different residues are found at the P4 to P4’ cleavage sites based upon the total number of each residue in tropoelastin isoform 2. (C) Bar chart of the absolute number of times different amino acids are located in the P1 or P1’ cleavage sites.

## METHODS

### Data Availability

All data used in the generation of this manuscript can be obtained by contacting the corresponding author. 3D printing files and instructions for device assembly can be obtained by contacting the corresponding author.

MatLab Code was modified from the referenced manuscript20 and can be downloaded from the supplementary material.

### Animals

#### Animal Use and Care

All animal use was approved by the Cincinnati Children’s Hospital Institutional Animal Use and Care Committee in accordance with the *Guide for the Care and Use of Laboratory Animals*. All animals used were on the C57BL/6 background and housed in a pathogen-free barrier facility given water and chow ad lib with 12-hour light-dark cycles. For each experiment, the number of male and female mice was roughly divided between experimental groups with animals of the same sex and experimental condition generally being kept in the same cage.

#### Animal Use

All animal use was approved by the Cincinnati Children’s Hospital Animal Use and Care Committee. Animals were housed in a barrier facility with twelve-hour light/dark cycles with food and water provided *ad lib*. C57BL/6 male and female mice were used for all experiments.

#### Cela1 Knockout Mouse

The guide RNAs indicated in Table S1 were validated in MK4 cells and subsequently co-injected with Cas9 mRNA into fifty single cell C57BL/6 embryos. Tail DNA was sequenced using exon 2 and 7 primers to identify founders.

#### Genotyping

A single mouse pup was found to have a heterozygous deletion in both exons 2 and 7. As the deletions were small, mice were genotyped using primers for exon 7 and digesting the PCR product with AluI (New England BioLabs) which differentially cut the PCR product.

#### Anti-AAT Oligo Administration

Anti-AAT or missense oligos at a dose of 100 mg/kg body weight were subcutaneously injected into wild type or *Cela1*^-/-^ mice weekly starting at 6 weeks of age. Lungs were collected for analysis after six weeks of therapy (five injections). The sequences of these oligos are proprietary (Ionis Pharmaceuticals).

#### Flexivent

Eight-week-old wild type and Cela1^-/-^ male and female mice were anesthetized and tracheally cannulated with a pre-calibrated 18 gauge flexivent cannula. Baseline lung mechanics were determined using a Flexivent FX using the manufacturer’s (Scireq) standard mouse baseline pulmonary dynamics program.

## Tissues

### Lung Tissue Processing

For histology, all lungs were inflated with 4% PFA in PBS at 25 cm H_2_O pressure, removed, and fixed overnight at 4°C. Lungs were dehydrated and paraffinized. The lung lobes were disarticulated and arranged to permit identification after sectioned. Five μm sections were cut from paraffin blocks and stained as below.

### Liver

Livers of oligo-treated mice were collected and stored at −80°C.

### Serum

At the time of sacrifice, blood from oligo-treated mice was centrifuged and serum stored at −80°C.

### Lung Wet to Dry Ratio

Postnatal day 14 (PND14) left lungs were weighed after animal sacrifice and allowed to desiccate overnight in a 60°C oven and weighed the next morning. Weight ratios were compared.

### Lung Homogenate Elastase Assay

Twenty μg of PND14 lung homogenate was incubated with 0.1 mg/mL Elastase Substrate (NS945, Elastin Products Company) and absorbance at 410 nm was assessed using a Spectramax plate reader.

## Molecular Biology Experiments on Tissues

### Western Blot

Western blots of mouse lung and pancreas homogenates, were performed using a previously validated anti-Cela1 guinea pig antibody^1^ and AlexaFlour680-conjugated, anti-tropoelastin antibody (AbCam, ab21600), and anti-β-actin antibody (Abcam, ab184092). Western blot of human serum specimens was performed using anti-Cela1 antibody (Abcam, ab118335) and anti-Serpina1 antibody (ab166610). Imaging and quantification was performed using a Licor Imaging System Image Studio Software (Licor).

### ELISA

Elisa for mouse Serpina1 protein was performed on serum using the Mouse Alpha 1 Antitrypsin ELISA kit (ab205088) from Abcam per manufacturer instructions.

### PCR

RNA was extracted from tissues lysates using the RNEasy Mini kit (Qiagen) and cDNA synthesized using High Capacity cDNA Reverse Transcriptase kit (Applied Biosystems). Taqman PCR was performed on RNA extracted from postnatal day 14 wild type and *Cela1*^-/-^, 8 week old untreated, control oligo-treated and anti-AAT oligo treated lungs using proprietary primers for mouse *tropoelastin*, *collagen1a*, *MMP14*, *emilin1*, *fibrillin1*, *fibulin1*, *neutrophil elastase*, *chymotrypsin-like elastase 1*, *tumor necrosis factor-α*, *interleukin-6*, *interleukin-10*, *interferon-α1/5/6*, *interferon-β*, *macrophage chemoattractant protein-1*, and *gapdh* (Applied Biosystems).

Since no taqman primer was validated for all five isoforms of murine Serpina1, we used Powerup SYBR Green (Applied Biosystems) and the *Gapdh* primers from the Rodent GAPDH Endogenous Control kit (Applied Biosystems) and forward and reverse primers that detected all five Serpina1 isoforms (Table S1)^17^. 77 (Serpina1) and 177 (GAPDH) base pair bands were confirmed by gel electrophoresis.

### Human Lung PCR

RNA was extracted from the lung of a young adult human motor vehicle accident victim. PCR for Cela1 was performed using Phusion DNA polymerase (New England BioLabs) and the primers listed in Table S1. Sequencing of PCR products was performed to verify sequence identity.

### Flow Cytometry

Using our previously established protocol^2^, from a single cell lung suspension lung cell type populations were identified by flow cytometry on a FACSCanto Cytometer (BD Biosciences) using Zombie Red (BioLegend, #423109), anti-vimentin-AF647 (Cell Signaling), anti-CD326/EpCAM-APC-eFlour780 (eBioscience #9856), anti-CD144/VE-cadherin-PE/Cy7 (BioLegend #138015), and anti-CD45-AF488 (Abd Serotec # MCA1031A488T). Appropriate secondary antibody conjugated OneComp eBeads (eBioscience) were used for compensation and fluorescence-minus-one samples were used to establish gates. Analysis was performed using FlowJo software.

## Imaging Experiments

### Mean Linear Intercept

H&E stained sections were imaged at 10X magnification collecting five images per lobe using a Nikon 90i upright microscope. Mean linear intercepts were determined by established methods^33,34^ and the average MLI per mouse used for statistical comparisons. For PND3 lung sections, 8 WT and 7 *Cela1*^-/-^ were analyzed. For PND14 6 WT, 5 *Cela1*^+/-^, and 9 *Cela1*^-/-^ were analyzed. For PN 8 weeks 8 WT and 8 *Cela1*^-/-^ were analyzed. RJ was not blinded to group allocation.

### Elastin Quantification

Lung sections were Hart stained by established methods^35^ and sections were tile scanned at 4X magnification. Using Nikon Elements software, lung area was determined by manually outlining each lobe. The tissue area was determined using an intensity mask. The total elastin area was determined using an intensity and hue mask.

### Measurement and Analysis of Airspace Diameters

Lungs were inflated, sectioned and stained with H&E. 4x tile scanned images were manually separated into each lobe using GNU Image Manipulation Program (GIMP, https://www.gimp.org/) and after filling in the background of segregated images, airspace sizes were determined after modifying a published MatLab program that was designed for quantifying the heterogeneity of airspace sizes in emphysema.^20^ After segmentation, major airways, vessels, and non-lung spaces were excluded, and airspace diameters for each lobe were calculated and then concatenated. Using Sigmaplot (Systat) histograms using 1,000 bins were created and the descriptive statistics of these histograms used for statistical comparisons.

### 3D-printed Confocal Microscope Stretching Device

A 3D-printed case with a site for a glass slide insert was constructed to house four servo motors which turned axles attached to Tyvek strips. The servo motors were controlled with a manual interface which incrementally turned the axle. The Tyvek strips were attached to 3D-printed clips which interfaced with a silicone mold with an aperture around which live lung sections were glued. This silicone mold was created by applying black silicone to a 3D-printed mold. To determine the change in aperture area with different steps area was measured. Device specifications and printing files are available through the corresponding author.

### Live Lung Sectioning and Securing to Silicone Mold

Lungs from 3 month old wild type and *Cela1*^-/-^ mice were inflated and sectioned with 8% gelatin in PBS as previously described^18^ except that instead of washing the sections with warm PBS after adhering to the silicone mold they were placed in DMEM media with 10% FBS, 1% penicillin/streptomycin, and 250 ng/mL amphotericin to remove the gelatin inflating media overnight. Stretch-inducible lung elastase activity was quantified as previously described.^18^

## Phylogeny Experiments

Ensembl protein and promoter sequences were downloaded between February 4-9, 2016 for all chordate sequences with annotations for Cela1, Cela2a, Cela2b, Cela3a, or Cela3b. The promoter sequences were limited to the upstream 200 base pairs. Phylogenetic trees were constructed with Megalign Pro (DNAStar) using Clustal W. Full length protein sequences were similarly processed. To determine the mammalian conservation of each isoform, we cross-checked Ensembl *Cela* protein sequences with Entrez Gene.

## Recombinant Protein Experiments

### Full length and truncated Cela1

Truncated (amino acids 27 to 266) murine Cela1 was cloned into the pET21a(+) plasmid with a 6-His Tag with sequence verification by Sanger sequencing. BL21(DE3) cells were transformed with this plasmid and recombinant protein purified on Ni-NTA columns (Qiagen). FLAG-tagged full length Cela1 was synthesized using a commercially available plasmid (Origene, MR203526) and transfected MK4 cells (derived from mouse metanephric mesenchyme)^36^ were lysed and purified plasmids were transfected into MK4 cells and cell lysates purified using magnetic anti-FLAG beads (Sigma). Western blot using a previously published guinea pig anti-Cela1 antibody^1^ demonstrated a band at the expected molecular weights for both proteins.

### Elastase Assay

The Enzcheck Elastase (Life Technologies) assay was used to incubate 1 μg of recombinant protein at 37°C for 4 hours with readings every ten minutes per manufacturer instructions. Results were analyzed as time vs. increase in fluorescent intensity of samples minus controls.

### Proteolysis of Elastin by Cela1 and Analysis of the Digests

Human tropoelastin (isoform 2) was recombinantly produced and mature, insoluble elastin was isolated from the foreskin of a five-year-old boy as described previously.^37^ Mature skin elastin and tropoelastin were dispersed or dissolved, respectively, at a concentration of 500 μg/mL in purified 1mg/mL Cela1 solution containing 10 mM Tris buffer, pH 7.5. All samples were digested for 24 h at 37°C. Proteolysis was stopped by adding trifluoroacetic acid to a final concentration of 0.5 % (V/V). Analysis of the enzymatic digests was carried out by nanoHPLC-nanoESI-QqTOF MS using an UltiMate 3000 RSLC nanoUHPLC system (Thermo Fisher) coupled to a QqTOF mass spectrometer Q-TOF-2 (Waters/Micromass, Manchester, UK) as described.^38^ Chromatographic separation of the peptides was performed at 40°C using a flow rate of 300 nL/min according to a protocol published earlier.^38^ Full scan MS data was acquired in the *m/z* range between 40 and 1550. Peptides were selected for tandem MS in data-directed acquisition mode using collision-induced dissociation by applying collision energies between 16 eV and 70 eV depending on charge state and *m/z*.

Product ion spectra were deconvoluted and deisotoped by MassLynx 4.1 with the add-on MaxEnt3 (Waters). Automated *de novo* sequencing followed by database matching was then performed using the software Peaks Studio (version 7.5; Bioinformatics Solutions, Waterloo, Canada).^39^ The search was restricted to the human section of the Swiss-Prot database extended by all splice variants of human tropoelastin. The enzyme was set to ‘none’ and mass error tolerances for precursor and fragment ions were set to 25 ppm and 0.1 Da, respectively. The peptide score threshold was decreased until a false discovery rate below 1% was achieved.

## Statistical Analysis

We estimated that 7-8 animals would be required per experimental group by performing a power analysis with a power of 90% to detect a 20% difference in means with a 10% standard deviation. No animals were excluded from analysis.

Sigmaplot (Systat) was used for all comparisons using t-test and one-way ANOVA for parametric values and Mann-Whitney U-test for nonparametric comparisons. P-values of 0.05 were considered statistically significant. Parametric data sets were displayed as bar graphs with error bars denoting standard error of the mean. Box plots were used for non-parametric values with boxes for 25^th^ and 75^th^ percentiles and whiskers indicating the 5^th^ and 95^th^ percentiles. For PCR experiments, some parametric data were displayed by box plot for clarity of presentation.

